# Divergence of variant binding/neutralizing antibodies following SARS-CoV-2 booster vaccines in myeloma: Impact of hybrid immunity

**DOI:** 10.1101/2023.08.17.553767

**Authors:** Alberto Moreno, Kelly Manning, Maryam I. Azeem, Ajay K. Nooka, Madison Ellis, Renee Julia Manalo, Jeffrey M. Switchenko, Bushra Wali, Jonathan L. Kaufman, Craig C. Hofmeister, Nisha S. Joseph, Sagar Lonial, Kavita M. Dhodapkar, Madhav V. Dhodapkar, Mehul S. Suthar

**Affiliations:** Emory Vaccine Center, Emory University, Atlanta, GA; Emory National Primate Research Center, Atlanta, GA; Division of Infectious Diseases, Department of Medicine, Emory University School of Medicine, Atlanta, GA; Department of Hematology/Medical Oncology, Emory University, Atlanta, GA; Aflac Cancer and Blood Disorders Center, Children’s Healthcare of Atlanta, Emory University, Atlanta, GA; Winship Cancer Institute, Atlanta, GA; Division of Infectious Diseases, Department of Pediatrics, Emory University School of Medicine, Atlanta, GA

## Abstract

We characterized virus-neutralization and spike-binding antibody profiles in myeloma patients following monovalent or bivalent-SARS-CoV-2 booster vaccination. Vaccination improves the breadth of binding antibodies but not neutralization activity against current variants. Hybrid immunity and immune imprinting impact vaccine-elicited immunity.

---

Hematological malignancies are associated with an increased risk of complications during SARS-CoV-2 infections (1, 2). Although vaccines have led to reduced risk of complications, vaccinated patients with hematologic malignancies still have a higher risk of hospitalization (3), as well as developing chronic SARS-CoV-2 infections (4-6), which have resulted in intrahost viral evolution and the emergence of variants of concern (7-9). Impaired induction of neutralizing antibodies (nAbs) against SARS-CoV-2 Omicron variants has been reported in myeloma (MM) patients following booster immunization with monovalent vaccines (10). Bivalent mRNA vaccines include both the ancestral WA1 and BA.5 spike. However, little is known about the breadth of nAbs induced by additional booster immunizations or immunization with monovalent or bivalent vaccines. In this study, we used an in vitro, live virus focus reduction neutralization test (FRNT) to determine the neutralization activity against WA1, BA.1, BA.5, BQ.1.1, and the contemporary XBB.1.5 Omicron subvariant in cohorts of MM patients that received monovalent or bivalent booster immunizations. We also evaluated spike-binding antibody responses against this panel of viruses using a Spike-specific electrochemiluminescence assay (11). All the participants provided written informed consent, and the Institutional Review Board of Emory University approved the study.

The study included two cohorts. The first cohort received anti-SARS-CoV-2 primary immunization and two monovalent booster immunizations, with the last monovalent booster administered 1-6 months after the second dose (n=101). The second cohort received the anti-SARS-CoV-2 primary immunization, two booster immunizations, and a bivalent booster administered 1-6 months after the third dose (n=42). The nAb titers against the ancestral strain (WA1) in patients that received the bivalent booster were 2.3-fold higher than neutralization titers in patients that received the monovalent booster (**Figure S1**). nAbs against BA.1 and BA.5 after monovalent or bivalent booster had lower titers compared to WA1, with a reduction in neutralization potency ranging between 7.1-8.2-fold for BA.1 and between 8.9-12.4-fold for BA.5 (**Figure S1**). Neutralizing antibodies against the newly circulating Omicron variants BQ.1.1 and XBB.1.5 were low or undetectable.

Detection of anti-nucleocapsid (NC) binding antibodies (900 AU/ml as cutoff) was utilized as evidence of recent SARS-CoV-2 exposure (12). In the monovalent-boosted group, 45.5% of the monovalent-boosted patients were anti-NC+ (n=46) (**Figure S2**). Next, we compared neutralization and antibody binding titers between individuals with low/undetectable anti-NC antibodies and those with recent SARS-CoV-2 exposure. In the monovalent-boosted group, the neutralization titers against WA1 were 11-fold higher in patients with recent SARS-CoV-2 exposure compared to unexposed patients (Geometric Mean Titer [GMT] 1379 vs 125). Neutralization titers against Omicron variants BA.1 (GMT 134 vs. 28), BA.5 (GMT 95 vs. 24), BQ.1.1 (GMT 38 vs. 20.5), and XBB.1.5 (GMT 30 vs. <20) were higher in SARS-CoV-2 exposed compared to unexposed patients (**Figure 1A**). We then assessed the temporal distribution of sampling among the monovalent-boosted group. In **Figure 1B**, antibody titers against WA1 are shown to indicate that most of the samples (69% and 59% in unexposed and exposed patients, respectively) were collected at time points under 50 days after immunization. However, a higher proportion of responders to multiple variants were observed among samples collected from exposed patients compared to unexposed patients at time points over 50 days after vaccination (**Figure 1C**).

**Figure 1.**
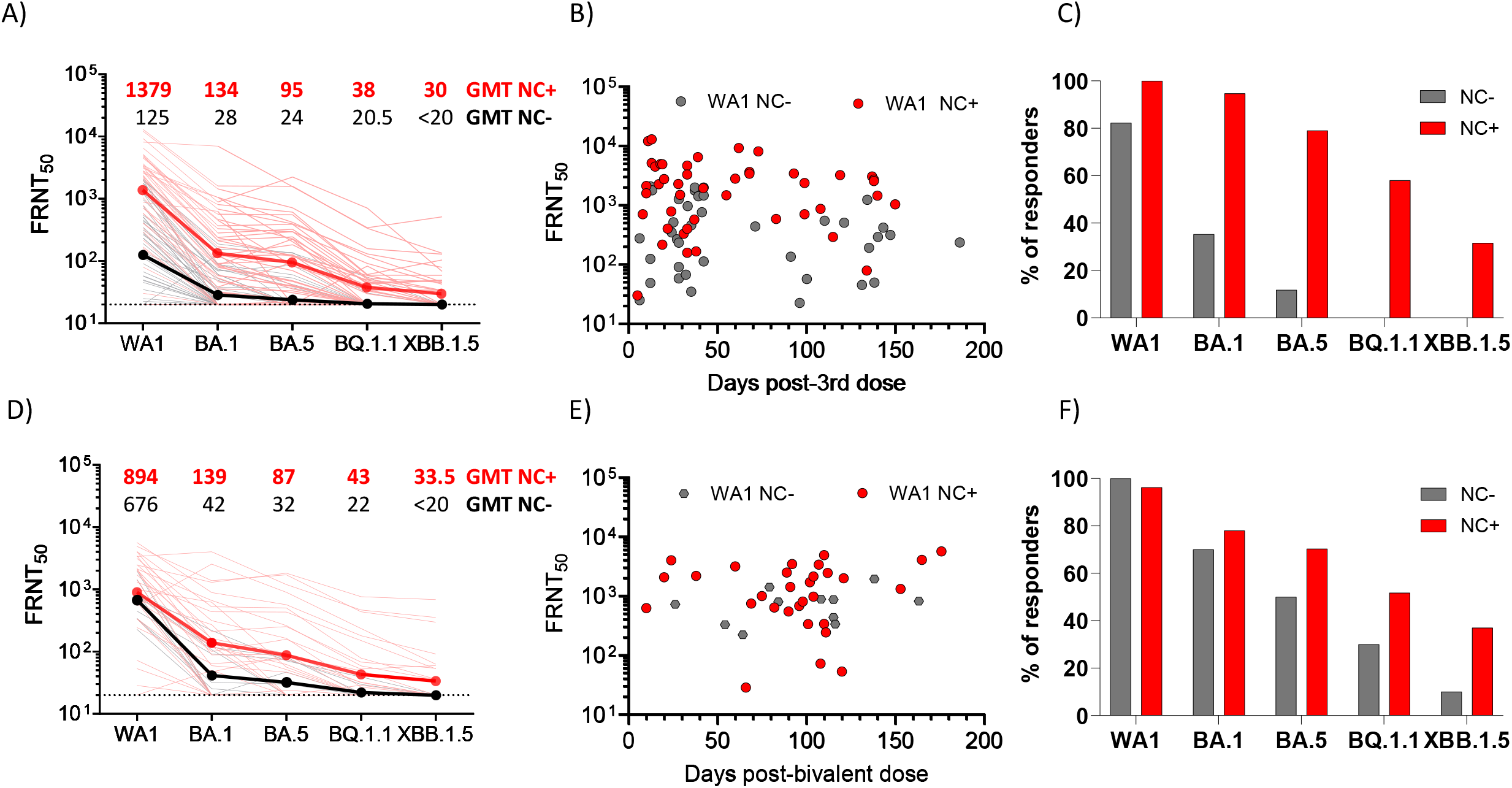
Live virus neutralizing antibody titers against WA1, BA.1, BA.5, BQ.1.1, and XBB.1.5 after monovalent or bivalent mRNA booster vaccination in patients with MM. (A) Comparative response of samples collected from MM patients after monovalent booster immunizations. Black lines, patients without evidence of natural exposure to SARS-CoV-2. Red lines, patients with natural exposure to SARS-CoV-2 by confirmation of positive anti-nucleocapsid binding antibodies. Dark black and dark red lines represent the geometric mean titers (GMT). The focus reduction neutralization test (FRNT50 [the reciprocal dilution of serum that neutralizes 50% of the input virus]) geometric mean titer (GMT) of neutralizing antibodies against the WA1 strain, and each Omicron subvariants is shown at the top of the panel, GMT values in black unexposed patients, GMT values in red patients exposed to SARS-CoV-2 infections. B) Time course of FRNT titers against WA1 in patients that received monovalent immunization and were unexposed (gray symbols) or exposed (red symbols) to SARS-CoV-2. C) Percentage of responders against individual variant tested at time points over 50 days, among unexposed (gray bars) or exposed (red bars) to SARS-CoV-2 that received monovalent booster immunization. D) Comparative response of samples collected from MM patients after bivalent booster immunizations. Black lines, patients without evidence of natural exposure to SARS-CoV-2. Red lines, patients with natural exposure to SARS-CoV-2 by confirmation of positive anti-nucleocapsid binding antibodies. Dark black and dark red lines represent the GMT of neutralizing antibodies against the virus tested. GMT of neutralizing antibodies is shown at the top of the panel, with GMT values in black unexposed patients and GMT values in red patients exposed to SARS-CoV-2 infections. E) Time course of FRNT titers against WA1 in patients that received bivalent immunization and were unexposed (gray symbols) or exposed (red symbols) to SARS-CoV-2. F) Percentage of responders against individual variant tested at time points over 50 days, among unexposed (gray bars) or exposed (red bars) to SARS-CoV-2 that received bivalent booster immunization.

Consistent with high community spread and breakthrough infections with the Omicron variant, we found that 73.8% of the bivalent-boosted patients were anti-NC+ (n=31) (**Figure S2**). In the bivalent-boosted group, the neutralization titers against WA1 showed modestly elevated titers (1.3 fold) in patients with recent SARS-CoV-2 exposure compared to unexposed patients (GMT 894 vs. 676). Neutralization titers were elevated against Omicron variants BA.1 (GMT 139 vs. 42), BA.5 (GMT 87 vs. 32), BQ.1.1 (GMT 43 vs. 22), and XBB.1.5 (GMT 33.5 vs. <20) in SARS-CoV-2-exposed compared to unexposed patients (**Figure 1D**). The temporal distribution of sampling among the bivalent-boosted group (**Figure 1E**) showed that most of the samples (91% and 87% in unexposed and exposed patients, respectively) were collected at time points over 50 days after immunization. A higher proportion of responders to multiple variants were present among samples collected from exposed patients compared to unexposed patients at time points over 50 days after vaccination (**Figure 1F**).

We next determined the IgG binding titers using Spike-specific electrochemiluminescence assays (11). Nearly all samples across the two cohorts tested positive for SARS-CoV-2 Spike-IgG binding (97%), except four unexposed SARS-CoV-2 patients receiving a monovalent booster. We tested a panel of pre-Omicron and Omicron Spike variants and found significantly higher binding titers in exposed SARS-CoV-2 patients that received a monovalent booster as compared to unexposed patients (P <0.0001; **Figure 2A**). In contrast, we did not observe a statistically significant difference in the bivalent vaccinated group (**Figure 2B**). In all the data points tested with sera samples collected from patients that received boosting immunization with monovalent or bivalent vaccines, no differences were observed between binding IgG titers to the Spike protein from early variants (B.1.1.7, B.1.351, and B.1.617.2) and the binding IgG titers to the Spike protein from the ancestral SARS-CoV-2 strain (**Figure S3**). However, in SARS-CoV-2 unexposed and exposed patients, differences were observed between the IgG antibody response against the Spike protein from the ancestral SARS-CoV-2 strain and the IgG response against most of the Omicron variants (**Figure S3**).

**Figure 2.**
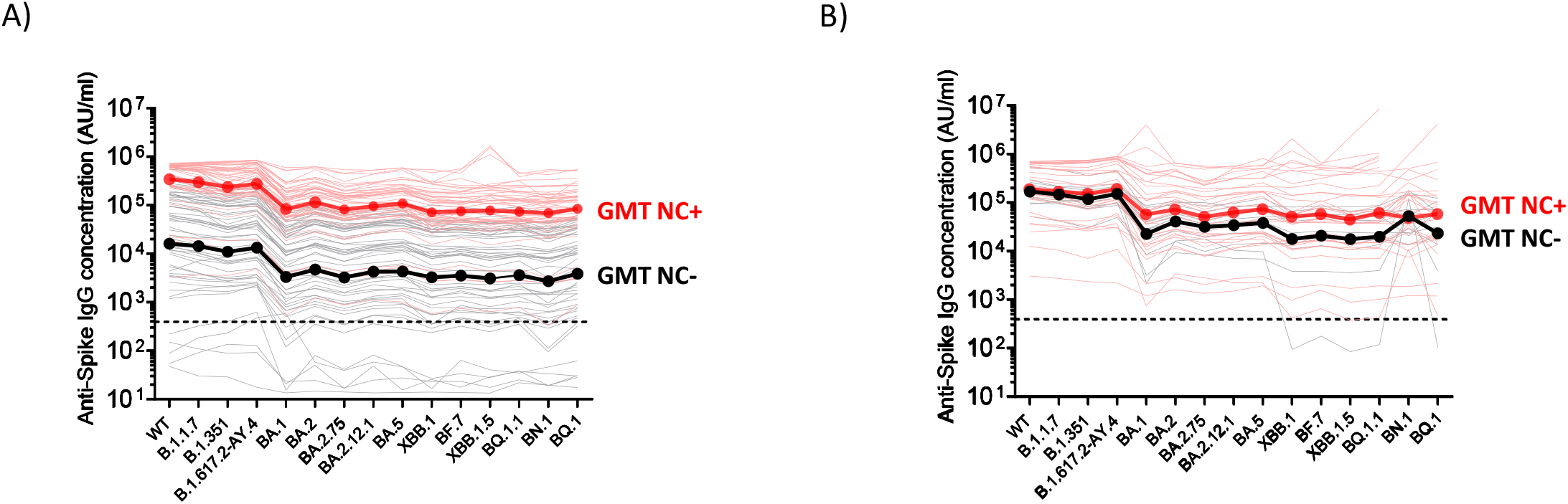
SARS-CoV-2 Spike-binding IgG antibody titers in MM patients that received a monovalent or bivalent booster immunization. Antibody responses were measured by electrochemiluminescence using the Meso Scale Discovery (MSD) platform. (A) Comparative response of samples collected from MM patients after monovalent booster immunizations. Black lines, patients without evidence of natural exposure to SARS-CoV-2. Red lines, patients with natural exposure to SARS-CoV-2 by confirmation of positive anti-nucleocapsid binding antibodies (titer >900 AU/ml). Dark black and dark red lines represent the geometric mean titers (GMT). B) Comparative response of samples collected from MM patients after bivalent booster immunizations. Black lines, patients without evidence of natural exposure to SARS-CoV-2. Red lines, patients with natural exposure to SARS-CoV-2 by confirmation of positive anti-nucleocapsid binding antibodies (titer >900 AU/ml). Dark black and dark red lines represent the geometric mean titers (GMT). Prepandemic plasma samples from healthy individuals were used to set the detection cutoff levels for SARS-CoV-2 Spike-specific IgG antibody titers.

We next analyzed the clinical correlates of neutralizing antibodies against WA1 and the contemporary XBB.1.5 variant as described (10, 13) (**Tables S1 and S2**). Anti-CD38 therapy correlated with reduced induction of nAb against WA1 in both cohorts of patients that received monovalent and bivalent boosters (**Table S2**). nAb against XBB-1 were low in all subsets and not impacted by these variables (**Table S1**).

This study shows that although bivalent vaccination induced significantly higher nAb titers against the ancestral strain in unexposed individuals, compared to those induced by monovalent boosters, titers against circulating Omicron variants (e.g., XBB1) were very low or undetectable. Hybrid immunity enabled higher induction of broadly reactive nAbs. However, there was no correlation between Spike-binding antibodies and neutralization against circulating variants suggesting that vaccination elicited antibodies that are not neutralizing. The inability to elicit antibodies against neutralizing epitopes suggests that a vaccination regimen with repeat boosters or using different vaccine platforms is needed to improve the effect of bivalent booster immunization in MM patients.

We have reported that a single monovalent booster vaccine is associated with increased nAb against the ancestral strain and the B.1.617.2 variant but not the Omicron variants in MM patients (10). It has been suggested that immune imprinting provided by prior infection or SARS-CoV-2 vaccination negatively impacts vaccine immunogenicity of booster immunizations (14). Consistent with this, we observed preferential boosting of nAb against the ancestral WA1 strain following booster immunization. Booster immunization particularly improved the breadth and longevity of the antibody response in patients with hybrid immunity. These results may be linked to the further expansion of class-switched memory B cells, described previously following monovalent boosters in MM (10). As with monovalent vaccines (10), patients with prior anti-CD38 antibodies also exhibit poor response to bivalent vaccines, which may relate to the adverse impact of these therapies on B/plasma cells and T-follicular-helper cells. The need for multiple vaccines in MM patients may not be SARS-CoV-2-specific, as multiple vaccines were also shown to improve seroprotection following influenza vaccination in a randomized trial in MM (15). Overall, our data show that most MM patients lack detectable nAb against current Omicron variants despite bivalent booster immunization. Spike-binding IgG antibodies against several Omicron variants were detected, suggesting that while immune imprinting may impact the induction of broadly neutralizing antibodies, reduced nAbs following vaccination in MM is not due to the inability to produce antibodies. High-risk cohorts of MM patients, such as those with prior CD38 or BCMA targeting therapies, may be at particular risk for ongoing reinfections with these viruses and may need consideration of newer approaches, including additional boosters or newer platforms.

## Acknowledgments

This work was supported in part by grants (NIH P51OD011132, 3U19AI057266-17S1, 1U54CA260563, HHSN272201400004C, NIH/NIAID CEIRR under contract 75N93021C00017 to Emory University) from the National Institute of Allergy and Infectious Diseases (NIAID), National Institutes of Health (NIH), Emory Executive Vice President for Health Affairs Synergy Fund award, the Pediatric Research Alliance Center for Childhood Infections and Vaccines and Children’s Healthcare of Atlanta, COVID-Catalyst-I^3^ Funds from the Woodruff Health Sciences Center and Emory School of Medicine, Woodruff Health Sciences Center 2020 COVID-19 CURE Award. This work was also supported in part by NCI U54 Seronet award CA260563. MVD is also partly supported by funds from the NIH R35CA197603 and the SCOR award from LLS. KMD is partly supported by funds from NIH CA238471 and AR077926, and LLS. Authors acknowledge the support of Cancer Tissue and Pathology, Data and Technology Applications, and Biostatistics shared resource of the Winship Cancer Institute of Emory University and NIH/NCI under award number P30CA138292. Funders played no role in the design and conduct of the study; collection, management, analysis, and interpretation of the data; preparation, review, or approval of the manuscript; and decision to submit the manuscript for publication.

## Authorship Contributions

A.M., Collected, analyzed, interpreted data, and wrote the original draft. K.M., M.I.A., M.E., and B.W., Performed assays. A.K.N., R.J.M., J.L.K., C.C.H., N.S.J., and S.L., Writing review and editing. J.M.S., Performed statistical analysis. K.M.D, Resources, formal analysis. M.V.D., and M.S.S., Conceptualization, resources, funding acquisition, writing review, and editing.

## Data Availability Statement

Data are available from the corresponding authors upon reasonable request.

