## Supplementary Material for "Divergence of variant binding/neutralizing antibodies following SARS-CoV-2 booster vaccines in myeloma: Impact of hybrid immunity"

### Supplementary Methods

*Patients & Patient Selection*

This study was approved by the Institutional Review Board of Emory University. Per protocol, patients were approached in myeloma clinics of Winship Cancer Institute without any selection bias. Patients who provided informed consent were eligible for a research blood draw. Blood samples were processed to isolate plasma and mononuclear cells.

*Nucleocapsid and Spike Binding Assay*

SARS-CoV-2 nucleocapsid and spike-specific IgG antibodies were detected using the Meso Scale Discovery (MSD) platform, V-PLEX SARS-CoV-2 Key Variant Spike Panel 1 Kit (catalog number K15651 (IgG)), and Panel 34 (catalog number K15690 (IgG)). Briefly, serum samples were diluted 1:5000 prior to the assay. Kit plates were blocked for 30 minutes using manufacturer-provided blocking buffer A. Blocking buffer was removed, and plates were washed 3X with wash buffer. Diluted serum samples, controls, and calibrators were added to the plates and incubated for 2 hours at room temperature in a plate shaker adjusted at 700 rpm. After a 2-hour incubation, plates were washed 3X, loaded with a solution containing MSD Sulfo-Tag anti-human IgG, and incubated for 1 hour. After incubation, the plates were washed 3X, and the detection read buffer was added immediately prior to the plate read. Raw data were parsed using Methodical Mind software.

*Viruses and cells*

VeroE6-TMPRSS2 cells were generated and cultured as previously described(1). nCoV/USA_WA1/2020 (WA1), closely resembling the original Wuhan strain, was propagated from an infectious SARS-CoV-2 clone as previously described(2). icSARS-CoV-2 was passed once to generate a working stock. The BA.1 isolate has been previously described(3). Omicron subvariants were isolated from residual nasal swabs: BA.5 isolate (EPI_ISL_13512579), provided by Dr. Richard Webby (St Jude Children's Research Hospital), BQ.1.1 isolate (EPI_ISL_15196219) and XBB.1.5 isolate (EPI_ISL_ 16818774) were provided by Dr. Benjamin Pinsky (Stanford University)  All variants were plaque purified and propagated once in VeroE6-TMPRSS2 cells to generate working stocks. All viruses used in this study were deep-sequenced and confirmed as previously described(4).

*Focus Reduction Neutralization Test*

FRNT assays were performed as described (1, 4, 5). In duplicate, serum samples were three-fold serially diluted down a plate, then mixed 1:1 with 100-200 PFU of WA, BA.1, BA.5, BQ.1.1 or XBB.1.5 (1:20 starting dilution with the virus). Dilution plates with virus-serum immune complex were incubated at 37°C for 1 hour. At 1 hour, the immune complex was loaded onto plates with a VeroE6-TMPRSS2 cell monolayer and incubated at 37°C for an additional hour. Following incubation, the immune complex was removed from cells and replaced with an 0.85% methylcellulose overlay. Cells were incubated at 37°C for variant-specific time intervals (16-40 hours). At the appropriate time points for each virus, cells were removed from the incubator, washed, and fixed with 2% paraformaldehyde. Cells were permeabilized and incubated with Alexa Fluor-647-conjugated SARS-CoV-2 (AF647-CR3022) for four hours at room temperature or 4°C overnight. After incubation, cells were washed, placed in Dulbecco's phosphate-buffered saline, and visualized on an ELISpot reader (CTL Analyzer).

*Statistical Analysis*

Antibody neutralization was quantified by counting the number of foci for each sample using the Viridot program(6). The neutralization titers were calculated as 1 - (ratio of the mean number of foci in the presence of sera and foci at the highest dilution of the respective sera sample). Each specimen was tested in duplicate. The FRNT50 titers were interpolated using a 4-parameter nonlinear regression in GraphPad Prism 9.4.1. Samples that do not neutralize at the limit of detection at 50% are plotted at 20 and used for geometric mean and fold-change calculations. The differences between all groups were determined with the Kruskal–Wallis test with Dunn's correction for multiple comparisons. Binomial logistic regression was used to identify the predictors for neutralizing antibody responses. Among variables statistically significant on univariate analysis, backwards elimination was utilized to identify relevant characteristics for inclusion in the final multivariable models, with an alpha value of 0.20. Unless otherwise noted, all statistical tests were two-sided, and statistical significance was assessed at the 0.05 level.

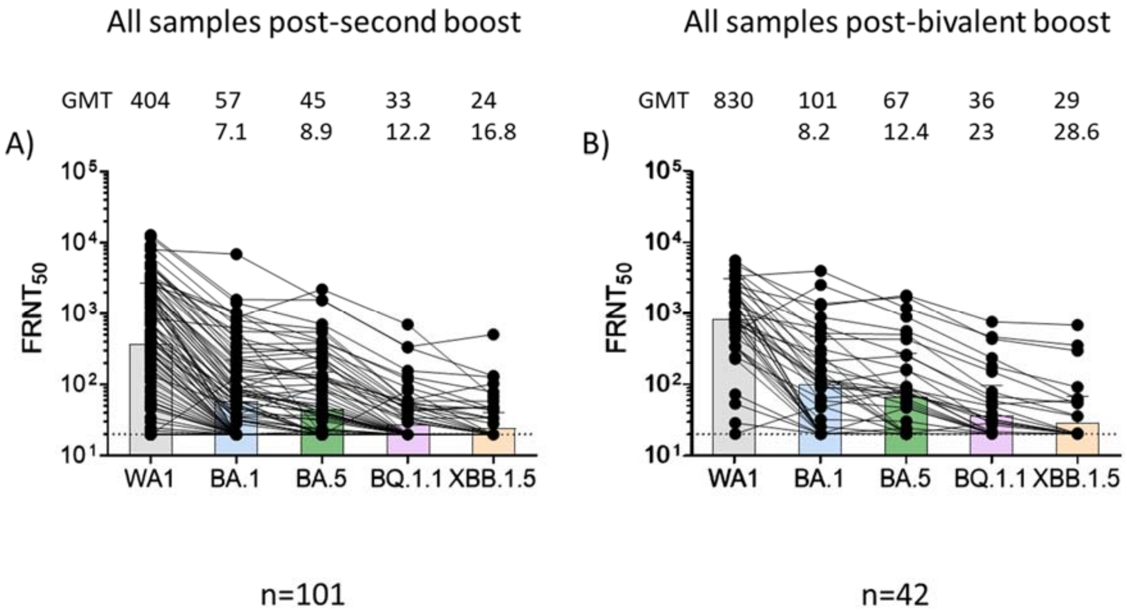

**Figure S1**. Neutralizing Responses against the WA1 Strain and Omicron Subvariants. Neutralization activity against the WA1 strain and the Omicron subvariants BA.1, BA.5, BQ.1.1, and XBB.1.5 in MM patients that received a monovalent booster (Panel A) or a bivalent booster immunization (Panel B). The FRNT50 geometric mean titer (GMT) of neutralizing antibodies against the WA1 strain and Omicron subvariants is shown at the top of each panel, along with the fold changes compared to WA1. The connecting lines between the variants represent matched serum samples. The horizontal dotted lines represent the limit of detection of the assay (FRNT50 GMT 20), and the colored bars the FRNT50 GMT.

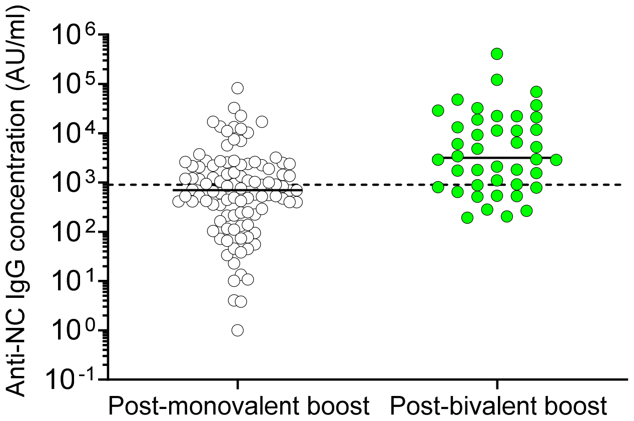

**Figure S2.** Natural exposure to SARS-CoV-2 infection was monitored by measuring nucleocapsid (NC)-specific IgG antibody titers by electrochemiluminescence assay. The horizontal dotted line represents the threshold for nucleocapsid positive signal (Anti-N IgG concentration =900 AU/ml)(7). N reactivity in the cohort of patients that received monovalent booster immunization was 45.5%, and in the cohort of patients that received bivalent booster immunization was 73.8%.

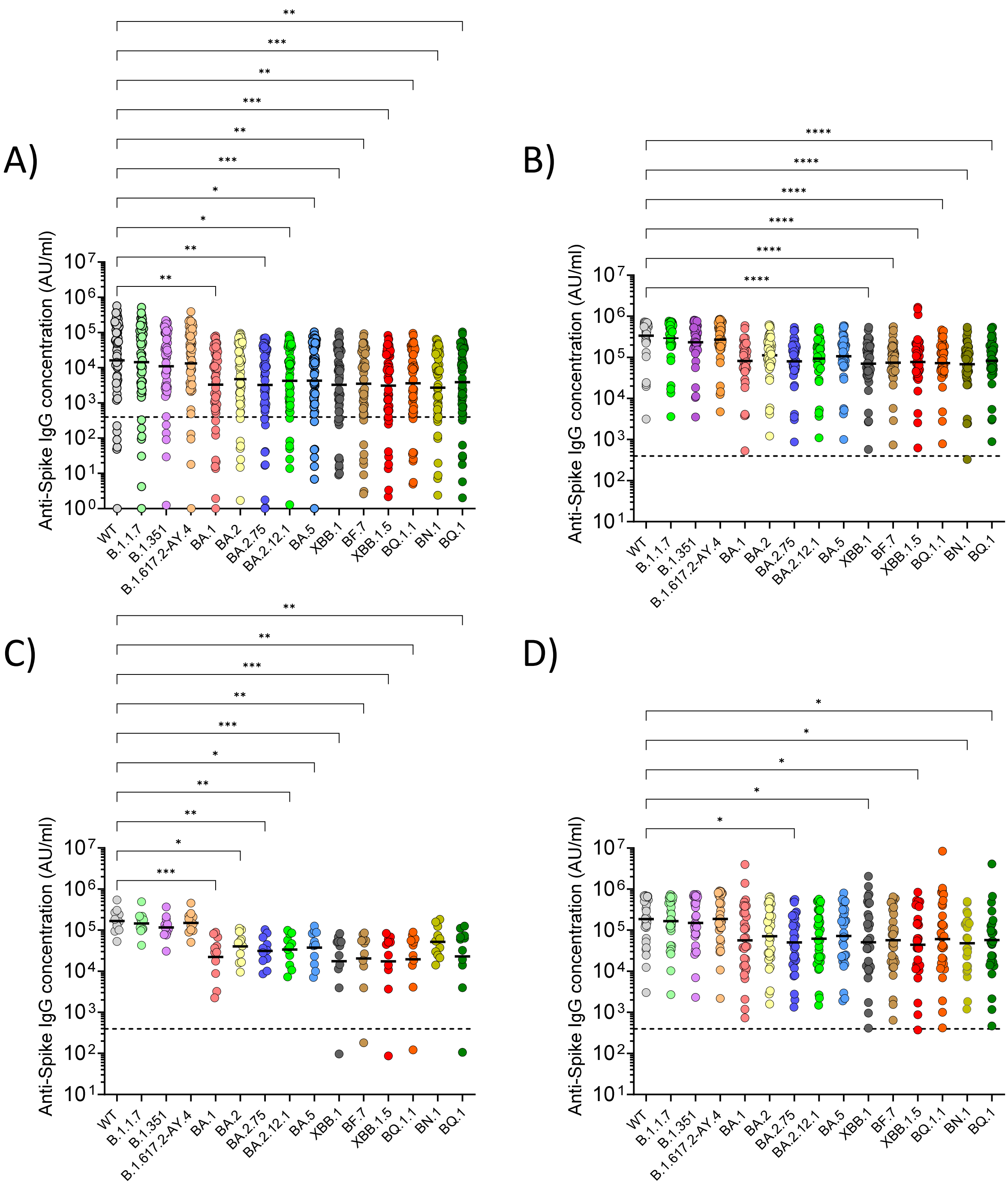
A)
B)
C)
D)
Anti-Spike IgG concentration (AU/ml)
WT
B.1.1.7
B.1.351
B.1.617.2-AY.4
BA.1
BA.2
BA.2.75
BA.2.12.1
BA.5
XBB.1
BF.7
XBB.1.5
BQ.1.1
BN.1
BQ.1

**Figure S3.** SARS-CoV-2 Spike-binding IgG antibody titers in MM patients that received a monovalent or bivalent booster immunization**.** A and B) Antibody titers after monovalent booster immunization that were previously unexposed to SARS-CoV-2 (Panel A) or previously exposed to SARS-CoV-2 (Panel B). C and D) Antibody titers after bivalent booster immunization that were previously unexposed to SARS-CoV-2 (Panel C) or previously exposed to SARS-CoV-2 (Panel D). Prepandemic plasma samples from healthy individuals were used to set the detection cutoff levels for SARS-CoV-2 Spike-specific IgG antibody titers. The differences between all groups were determined with the Kruskal–Wallis test with Dunn's correction for multiple comparisons. * $p \leq 0.05$, ** $p \leq 0.01$, *** $p \leq 0.001$, **** $p \leq 0.0001$.

**Table S1a: Univariate logistic regression – XBB.1.5 - Monovalent**

| | | | XBB_1_5=Positive | |
|---|---|---|---|---|
| Covariate | Level | N | Odds Ratio (95% CI) | OR P-value |
| Race | Black | 39 | 2.56 (0.88-7.45) | 0.084 |
| | Other | 59 | - | - |
| Sex | Female | 47 | 0.39 (0.12-1.20) | 0.100 |
| | Male | 51 | - | - |
| Age <=65 | Yes | 31 | 1.63 (0.56-4.80) | 0.372 |
| | No | 66 | - | - |
| Prior LOT (>2) | Yes | 32 | 1.40 (0.48-4.12) | 0.541 |
| | No | 60 | - | - |
| IgG <=400 | Yes | 21 | 1.16 (0.33-4.01) | 0.816 |
| | No | 77 | - | - |
| antiCD38 | Yes | 20 | 0.20 (0.03-1.64) | 0.135 |
| | No | 78 | - | - |
| Len maintenance | Yes | 28 | 1.46 (0.48-4.43) | 0.501 |
| | No | 70 | - | - |
| antiBCMA | Yes | 3 | - | - |
| | No | 95 | - | - |
| Prior SARS CoV-2 exposure | Positive | 44 | - | - |
| | Negative | 54 | - | - |

**Table S1b: - Univariate logistic regression – XBB.1.5 - Bivalent**

|  |  |  | XBB_1_5=Positive<br>---------------------------------------<br>- |  |
|---|---|---|---|---|
| **Covariate** | **Level** | **N** | **Odds Ratio (95% CI)** | **OR P-value** |
| Race | Black | 19 | 1.20 (0.29-4.94) | 0.800 |
| | Other | 18 | - | - |
| Sex | Female | 20 | 1.11 (0.27-4.55) | 0.880 |
| | Male | 18 | - | - |
| Age <=65 | Yes | 8 | 3.29 (0.65-16.67) | 0.151 |
| | No | 30 | - | - |
| Prior LOT (>2) | Yes | 9 | 0.60 (0.10-3.51) | 0.574 |
| | No | 28 | - | - |
| IgG <=400 | Yes | 10 | 1.07 (0.22-5.21) | 0.932 |
| | No | 28 | - | - |
| antiCD38 | Yes | 13 | 0.32 (0.06-1.79) | 0.196 |
| | No | 25 | - | - |
| Len maintenance | Yes | 7 | 4.57 (0.82-25.46) | 0.083 |
| | No | 31 | - | - |
| antiBCMA | Yes | 2 | - | - |
| | No | 36 | - | - |
| Prior SARS CoV-2 exposure | Positive | 28 | 5.00 (0.55-45.39) | 0.153 |
| | Negative | 10 | - | - |

**Table S2a: - Univariate linear regression - WA1 - Monovalent**

| | | | Log WA1 | |
|---|---|---|---|---|
| Covariate | Level | N | B (95% CI) | B P-value |
| Race | Black | 39 | 0.17 (-0.63-0.97) | 0.670 |
| | Other | 59 | - | - |
| Sex | Female | 47 | -0.44 (-1.22-0.34) | 0.272 |
| | Male | 51 | - | - |
| Age <=65 | Yes | 31 | 0.76 (-0.08-1.59) | 0.076 |
| | No | 66 | - | - |
| Prior LOT (>2) | Yes | 32 | -0.74 (-1.58-0.11) | 0.088 |
| | No | 60 | - | - |
| IgG <=400 | Yes | 21 | -0.19 (-1.14-0.77) | 0.703 |
| | No | 77 | - | - |
| antiCD38 | Yes | 20 | -0.98 (-1.93--0.03) | **0.044** |
| | No | 78 | - | - |
| Len maintenance | Yes | 28 | 0.84 (-0.01-1.69) | 0.054 |
| | No | 70 | - | - |
| antiBCMA | Yes | 3 | -3.03 (-5.23--0.83) | **0.007** |
| | No | 95 | - | - |
| Prior SARS CoV-2 exposure | Positive | 44 | 2.39 (1.76-3.02) | **<.001** |
| | Negative | 54 | - | - |

**Table S2b: - Univariate linear regression - WA1 - Bivalent**

| | | | Log WA1 | |
|---|---|---|---|---|
| Covariate | Level | N | B (95% CI) | B P-value |
| Race | Black | 19 | -0.48 (-1.35-0.38) | 0.271 |
| | Other | 18 | - | - |
| Sex | Female | 20 | 0.26 (-0.59-1.11) | 0.545 |
| | Male | 18 | - | - |
| Age <=65 | Yes | 8 | 0.67 (-0.36-1.69) | 0.202 |
| | No | 30 | - | - |
| Prior LOT (>2) | Yes | 9 | -0.61 (-1.60-0.38) | 0.225 |
| | No | 28 | - | - |
| IgG <=400 | Yes | 10 | 0.26 (-0.70-1.23) | 0.595 |
| | No | 28 | - | - |
| antiCD38 | Yes | 13 | -0.87 (-1.72--0.01) | **0.048** |
| | No | 25 | - | - |
| Len maintenance | Yes | 7 | 0.71 (-0.37-1.79) | 0.198 |
| | No | 31 | - | - |
| antiBCMA | Yes | 2 | 0.67 (-1.23-2.57) | 0.488 |
| | No | 36 | - | - |
| Prior SARS CoV-2 exposure | Positive | 28 | 0.22 (-0.75-1.18) | 0.661 |
| | Negative | 10 | - | - |

7. Chang A, Akhtar A, Lai L, Orellana-Noia VM, Linderman SL, McCook-Veal AA, et al. Antibody binding and neutralization of live SARS-CoV-2 variants including BA.4/5 following

booster vaccination of patients with B-cell malignancies. Cancer Res Commun. 2022;2(12):1684-92.
